# Parameter scaling of multivariate Granger causality

**DOI:** 10.1101/2025.10.01.679714

**Authors:** Thomas Pirenne, Esther Florin

## Abstract

Estimating causal interactions between signals provides unique insights into their dynamics, and causal inference has been widely applied to electrophysiological data to elucidate brain communication. Multivariate autoregressive models (MVAR) form the basis of most causal estimation methods. However, the high dimensionality of whole-brain data renders MVARs difficult to estimate reliably, and reducing the dimensions to a reasonable range affects causal inference. To address these scalability limitations, we develop sparse Multivariate Granger Causality (sMVGC), a novel method premised on the assumption that true causal connections between signals are sparse, thereby constraining the candidate search space and improving scalability. To motivate sMVGC empirically, we simulate electrophysiological data with known causalities and model how the number of samples, signals, and MVAR order affect the performance and computation time of current algorithms. All algorithms scale at least quadratically with these parameters, yet differ in their sensitivity to signal versus sample count, and the sample requirements for accurate inference scale with the number of signals. Guided by these findings, sMVGC improves parameter scalability while preserving estimation accuracy, and we provide practical parameter ranges and model selection guidance for real-world analyses.

## 1 Introduction

Estimating causal interactions between signals provides unique insights into their underlying dynamics. In neuroscience, causal interactions between different brain regions are commonly estimated to shed light on their functional organization. Many causality inference methods exist, including Granger causality based on regression [1], information-theoretic approaches [2], or causal network learning algorithms [3–5]. Among these, Granger causality analyses based on multivariate autoregressive models (MVAR) are distinctive in yielding conditional causality measures, i.e., they explicitly account for mediating effects between all signals [6, 7]. Their central assumption is that a first signal Granger causes a second if the information carried by the first signal improves the forecasting of the second [8]. Multivariate Granger causality (**MVAR**) [1] and (generalized) Partial Directed Coherence (**PDC**) [9] are two common methods to derive causality measures from MVAR parameters.

Three parameters determine the scale of MVAR models in causal estimation: the number of signals involved (number of channels, *nc*), the length of their recorded activity (number of samples, *ns*), and the model order (order, *no*). Failing to consider all relevant sources can lead to wrong causal estimation [10–12]. While considering interactions between all involved signals limits detection errors, it also increases the number of signals between which causality is estimated resulting in a quadratic rise in its search space (curse of dimensionality). It is widely accepted that causality estimation methods have an upper bound in the number of signals between which it can be estimated reliably [7, 12]. In addition, estimating causality between more signals requires longer recordings, further increasing the scale of the data [10, 12]. Finally, the order of causality estimation, i.e., how far back in time to look for causality between signals, further increases the dimensionality of MVAR parameters.

The investigation of electrophysiological brain activity usually requires limiting these three parameters, affecting the quality of causal estimations. The number of considered signals is often reduced to a few regions of interest [13–17], effectively limiting the number of signals, to reduce computational costs. This requires dimensionality reduction of the data, which affects causal estimation [18]. Investigating causal dynamics at a timescale unadapted to the sampling rate of data requires either downsampling the data or increasing the model order. However, the curse of dimensionality limits the model order, and downsampling beyond causal interaction delay was shown to create spurious and undetected causalities [19, 20].

MVARs can be estimated by many algorithms. Here we consider the *Levinson-Wiggins-Robinson* regression (**LWR**) [1, 21, 22], which is common in neuroscience, and the classic *ordinary least-squares regression* (**OLS**), which was shown to outperform LWR [23]. More recent methods to estimate the MVAR parameters try to reduce the minimum data length required to estimate causality for a fixed number of sources and in different noise conditions. For instance, Antonacci et al. extended OLS with a *L*_2_ penalized regression of the least absolute shrinkage and selection operator (**LASSO**) to provide accurate estimations from shorter segments [24]. Combining artificial neural networks with state-space models yields more accurate causal predictions in short segments and sparse conditions [25]. Around the same time, Li et al. proposed a Bayesian learning method based on Sparse Bayesian Learning (**SBL**) to estimate MVAR parameters [26]. It was designed to estimate causality more robustly in short segments and low signal-to-noise ratio (**SNR**) conditions. More recently, Liu et al. built upon this method by having the fitting error follow a Laplace distribution, making it more robust to outliers [17].

While the curse of dimensionality in causal inference is commonly acknowledged, the exact limits are unknown. In this work, we investigate the effect of the main parameters driving the dimensionality of causal inference (*nc, ns, no*) on the performance and computation time of the most common MVAR-based algorithms. Our analyses clearly identify the dimensionality limits of current causal inference algorithms. To address these limits, we propose sparse Multivariate Granger Causality (**sMVGC**), a variation of MVGC [1] that restricts the search space to a tractable subset of candidate connections. Under the assumption of sparse connectivity, sMVGC enables conditional Granger causality estimation across substantially larger numbers of signals.

## 2 Methods

As illustrated in figure 1, our investigation is structured in three components: causality simulation, estimation, and scalability analysis. All analyses depicted were performed without added external noise. They were run in parallel with a signal-to-noise ratio (SNR) of 1. The analyses highlight the scalability limitations of existing methods, inspiring an alternative approach designed to overcome them: sparse-MVGC.

**Figure 1:**
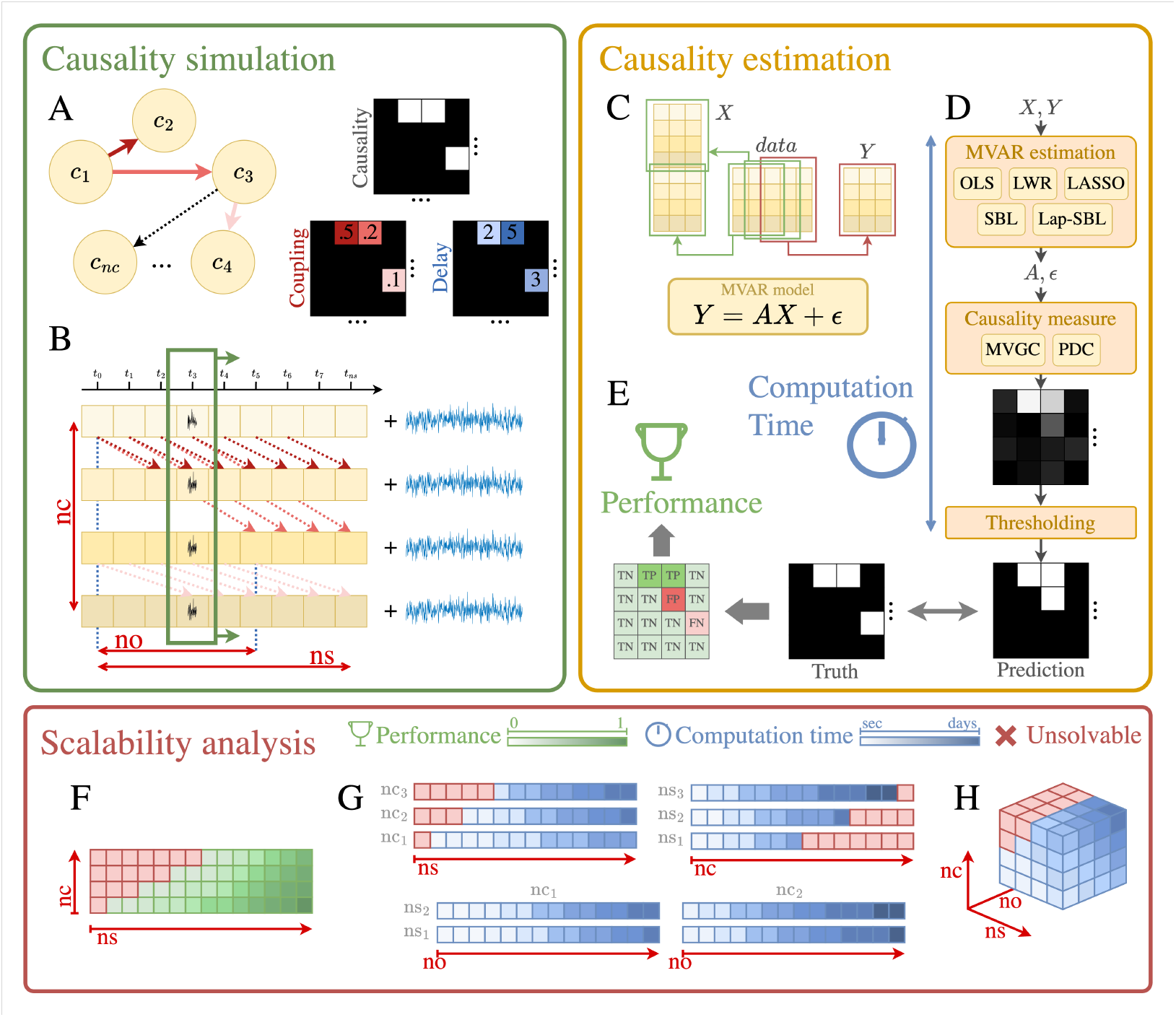
**A.**Causality network formation. **B**. Data generation. **C**. Multivariate autoregressive model. **D**. Causality estimation. **E**. Performance measures of causality estimation. **F**. Prediction performance with respect to *ns* and *nc*. **G**. Computation time with respect to *ns, nc, no* independently. **H**.Computation time with respect to *ns, nc*, and *no* interdependently.

### 2.1 Causality Simulation

Causally interconnected data is simulated with multiple combinations of parameters *nc, ns*, and *no*.

#### Causality network formation

Simulating a causal network of *nc* signals requires three *nc × nc* matrices: *causality* (*C*), *coupling* (*S*), *delay* (*D*). The binary matrix *C* encodes the ground-truth causal structure, where *C*_*ij*_ = 1 indicates that signal *i* causes signal *j*. Causal connections are assigned randomly in the upper triangular portion of *C* to prevent feedback loops, which introduce nonlinearities that can destabilize signals. Network density is set such that average node degree equals 2 for *nc ≥* 8, and the connection proportion is set to 0.25 for *nc <* 8. While the simulated signals are meant to fit the reality of causal relationships in natural signals, some modeling choices remain arbitrary. In supplementary experiments, we additionally evaluate denser networks with average node degrees of 10 and 50. Feedback loops are addressed separately by mirroring *C* (i.e., *C*_*feedback*_ = *C* + *C*^*T*^), rendering all connections bidirectional. The coupling matrix *S* assigns a random strength *U ∈* [*−*1.0, 1.0] to each causal connection. A negative coupling strength indicates a negative relationship between cause and effect, i.e. an amplitude increase in the cause is associated with an amplitude decrease in the consequence. Assigned coupling strengths (*S*^*ass*^) are normalized to sum to at most 1 per consequence. In supplementary analyses, we further scale down *S*^*ass*^ to match the external noise proportion (*n*_*p*_), yielding effective coupling strengths (*S*^*eff*^) as defined in equation (3). The delay matrix *D* assigns a random integer delay in [1, *no*_*data*_] to each causal connection, representing the sample lag between cause and effect, as defined in equation (4).

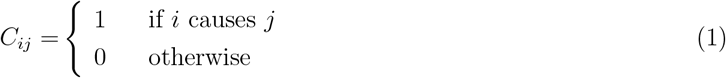

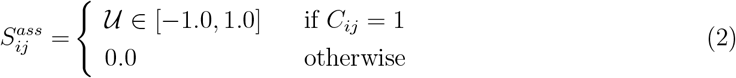

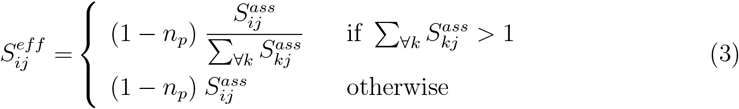

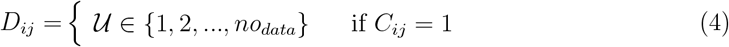

#### Causality data generation

Causal time series are generated iteratively from the causality, coupling and delay matrices. External noise, scaled by proportion *n*_*p*_ *∈* [0, 1], is added to each time series upon completion, where *n*_*p*_ = 0.0 denotes no noise and *n*_*p*_ = 1.0 denotes noise only. The corresponding signal-to-noise ratio (SNR) is given by:

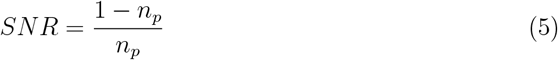

The primary parameters of interest in the analysis are *nc* (number of signals), *ns* (number of samples), and *no*_*data*_ (order, i.e., largest generated delay). Linear causality of delay *d* between *X → Y* is simulated as:

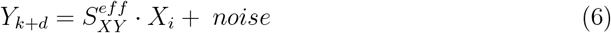

In supplementary analyses, we extend this framework in two ways. First, we substitute white external noise with *pink* noise (power decreasing by 3dB per octave), which may better approximate the noise structure of electrophysiological signals. Second, we assess the capacity of MVAR-based estimation to detect non-linear causalities. Specifically, we simulate quadratic and cosine causalities, defined respectively as:

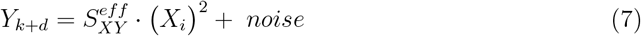

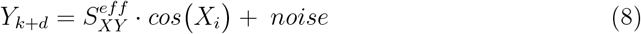

### 2.2 Causality Estimation

For each set of simulated data, with each algorithm of interest, causality is estimated.

#### MVAR modeling

Causality estimation begins by fitting an MVAR model to the data. The model uses the first *no* lags of all *nc* signals (*X*: (*nc · no*) *×* (*ns − no*)) to forecast their last *ns − no* samples (*Y* : (*nc*) *×* (*ns − no*)), by solving:

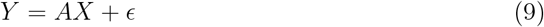

where *A* (*nc ×* (*nc · no*)) is the parameter matrix and *ϵ* ((*nc*) *×* (*ns −no*)) the residual. In our main experiments, the model order is assumed to match the underlying causal delay of the data (*no* = *no*_*data*_). While ways to estimate the model order have been investigated in previous work [1], in our supplementary experiments, we consider cases of both under- and overestimated orders.

Five algorithms for estimating *A* are evaluated. *Ordinary least squares* (**OLS**) regression minimizes the residual *ϵ* [8]. The *Levinson, Wiggins, Robinson* algorithm (**LWR**) reformulates the MVAR as Yule-Walker equations and solves them recursively via maximum entropy [27]. The *Least Absolute Shrinkage and Selection Operator* (**LASSO**) extends OLS with an L1 penalty to exploit causal sparsity [13, 28], with the regularization parameter *λ* optimized via generalized cross-validation. *Sparse Bayesian learning* (**SBL**) employs Bayesian inference with an expectation-maximization (**EM**) algorithm, assuming Gaussian residuals; an uninformed prior is used here given the absence of domain-specific prior knowledge [26]. Finally, *Laplace-sparse Bayesian Learning* (**Lap-SBL**) improves robustness to outliers by replacing the Gaussian residual assumption with a multivariate Laplace distribution [17].

#### Causality metric

Estimating *A* does not directly yield causal predictions. Causality must be inferred from either *A* or the residuals *ϵ. Multivariate Granger causality* (**MVGC**) [1] follows the latter approach, fitting one *full* regression (*A*^*f*^, *ϵ*^*f*^) and *nc reduced* regression, each excluding signal *j* (*A*^*rj*^, *ϵ*^*rj*^). The causal effect of *j* on *i* is then estimated as the log-ratio of reconstruction error covariances:

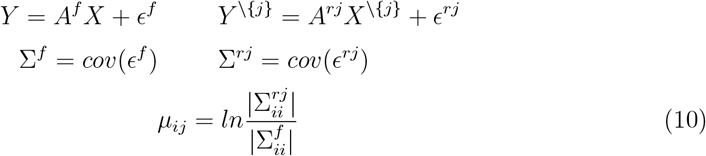

In contrast, *partial directed coherence* (**PDC**) [9, 29] derives causality directly from *A*, expressing the relative contribution of *j*’s past to forecasting *i* in the frequency domain:

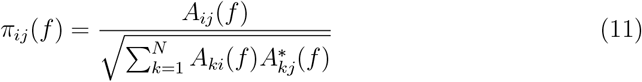

where Σ_*i*_ |*π*_*ij*_ (*f*)|^2^ = 1. To address sensitivity to signal scaling, we adopt the *generalized* PDC formulation [30]:

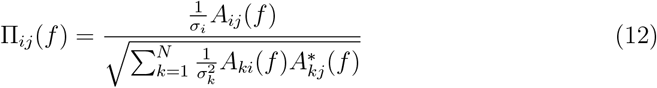

where 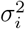 is the variance of *ϵ*_*i*_. All subsequent references to PDC denote this generalized version. Both methods incorporate significance tests (F-test and likelihood ratio test, respectively) to reject spurious connections [1, 30], with thresholds corrected for multiple comparisons as *p <* (0.001 *× nc*^2^).

#### Performance measures

Computation time is recorded for the full causality estimation pipeline across all algorithms, run on identical hardware and software. Since our analyses target relative scalability rather than absolute timing, the specific hardware matters less than that the computations were run in similar conditions. Estimated causality is then compared against the ground truth, yielding the standard confusion matrix counts: true positives (TP), true negatives (TN), false positives (FP), and false negatives (FN). Given the inherent sparsity of causality networks, particularly as *nc* increases, performance is evaluated using the f1-score, which balances precision and recall into a single metric:

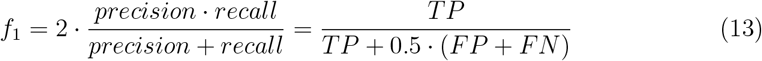

### 2.3 Scalability Analysis

In our analyses, we simulate and estimate causality (figure 1 **A-E**) with varying combinations of *nc, ns, no*. For each combination, we measure the computation time and evaluate the prediction performance as described above. We combine our base causality measure with each of the MVAR estimation methods to investigate the effect of the latter: *MVGC-LWR, MVGC-OLS, MVGC-LASSO, MVGC-SBL, MVGC-lapSBL*. To investigate the effect of the causality measure, we combine PDC with our base MVAR estimation methods: *PDC-LWR, PDC-OLS*. Causal inference of datasets generated with different sets of parameter combinations (*nc, ns, no*) provides distinct insights into the methods’ scalability. In all parts of these experiments, we stop increasing the parameter values (*nc, ns, no*) when computation time increases above 1000*s*. This makes our experiments tractable in time and is high enough to capture the nature of the time growth, e.g., linear, quadratic, or cubic.

The first set of combinations of interest is *no* = 10, *nc ∈* [10; 480], *ns ∈* [100; 10, 000], in steps of 10 and 100 respectively (figure 2, 3A). Causal inference depends on *ns*. The f1-scores of this set of combinations are expected to reveal to what extent *nc* affects this dependence. A special focus is given on the smallest *ns* which yields an *f* 1*score >* 0 (*ns*\*). This represents the minimal amount of data required to start inferring causality with some accuracy. From the same set of combinations, we identify this *ns*\* with respect to *nc* (figure 3B). This allows comparing inference methods in their ability to predict causality from short segments.

**Figure 2:**
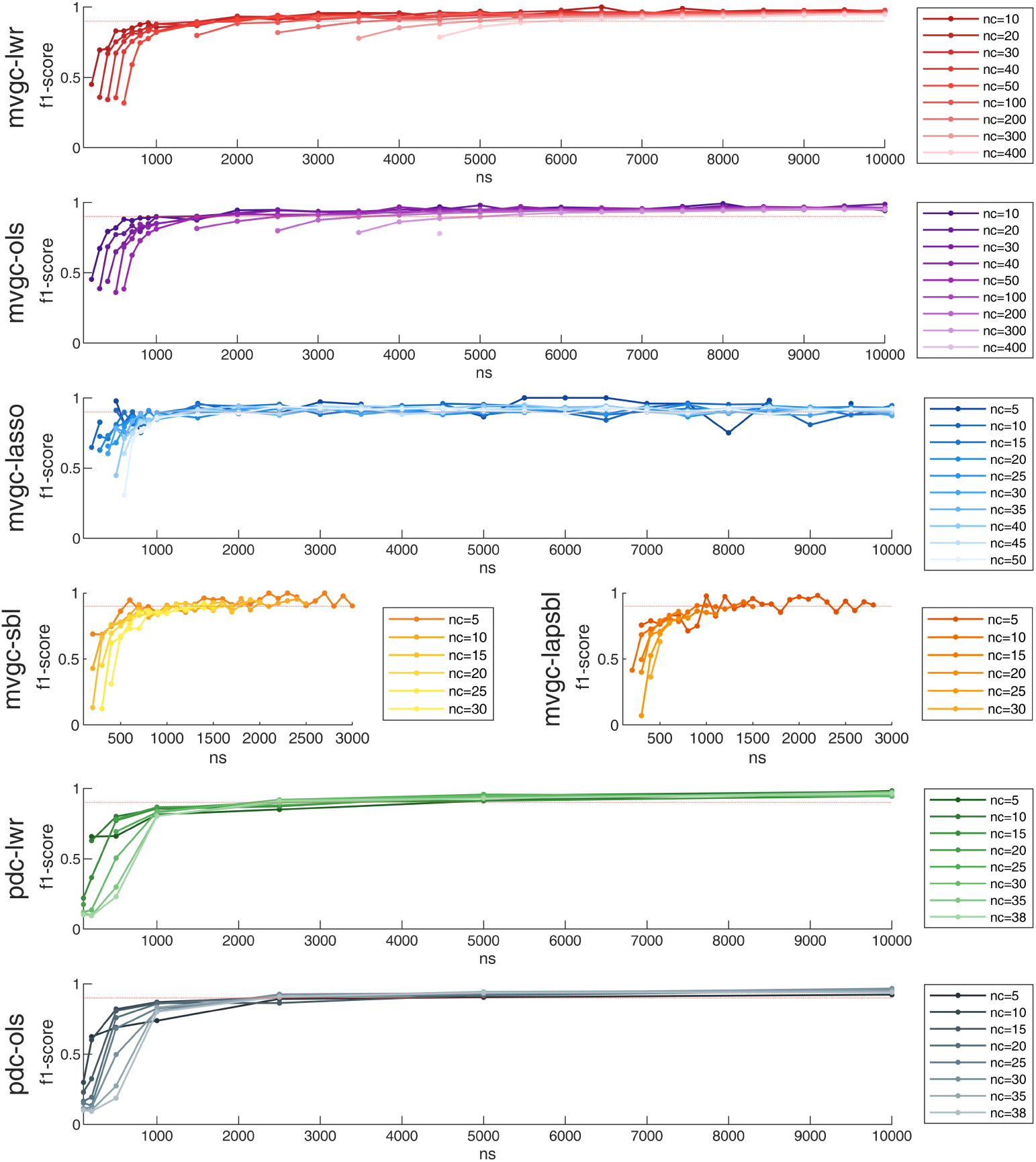
F1-scores (y-axis) obtained by the methods of interest while predicting causality from data including different amounts of signals *nc* (dark-to-light), with increasing amounts of samples *ns* (x-axis).

**Figure 3:**
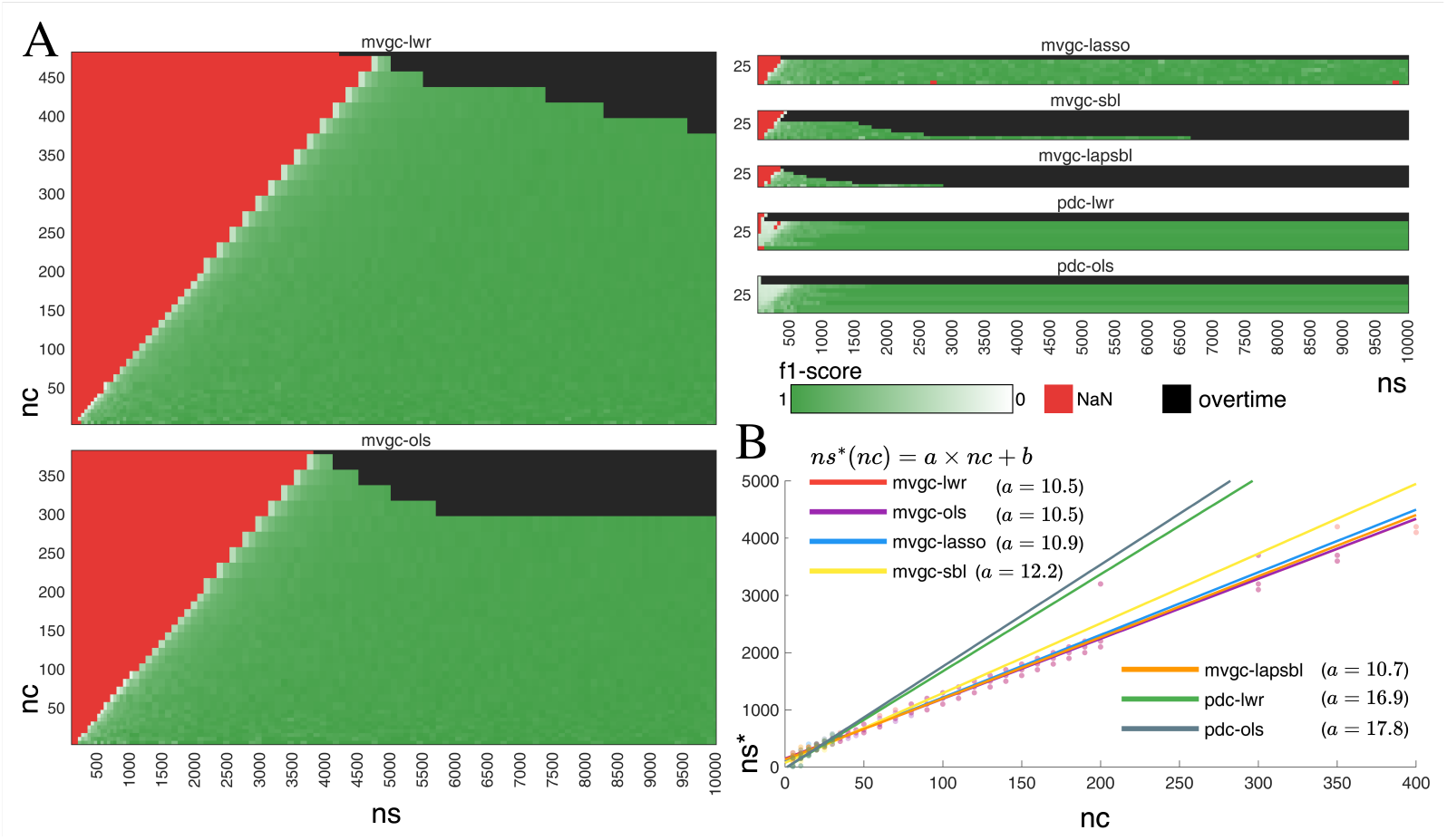
**A**: Feasibility area of each method of interest. Red: the method failed to yield a prediction, or the prediction obtained *f* 1*s* = 0. Light-dark green: the method obtained *f* 1*s >* 0 in reasonable time. Black: the method needs *time >* 1000*s* to potentially yield a prediction. **B**: Linearly regressed *ns*\*(*nc*), i.e., such that, for a given *nc, ns < ns*\* yields *f* 1*s* < 0.5 and *ns > ns*\* yields *f* 1*s >* 0.5.

The second set of combinations of interest is *ns ∈* [50; 10, 000], *nc ∈* [5; 400], *no ∈* [5; 200] in steps of 50, 5, and 5 respectively. The analysis of this set investigates how *ns, nc*, and *no* affect computation time independently. For each parameter, we display the evolution of its computation time through its whole range, while fixing the other two at meaningful values. We visualize the effect of *ns* by fixing *no* = 10 and *nc ∈* {10, 30, 200} (figure 4A). The effect of *nc* is depicted by fixing *no* = 10 and *ns ∈* {1000, 2000, 10000} (figure 4B). The effect of *no* is shown by fixing *nc ∈* {10, 30, 200} and *ns ∈* {1000, 10000}) (figure 4C).

**Figure 4:**
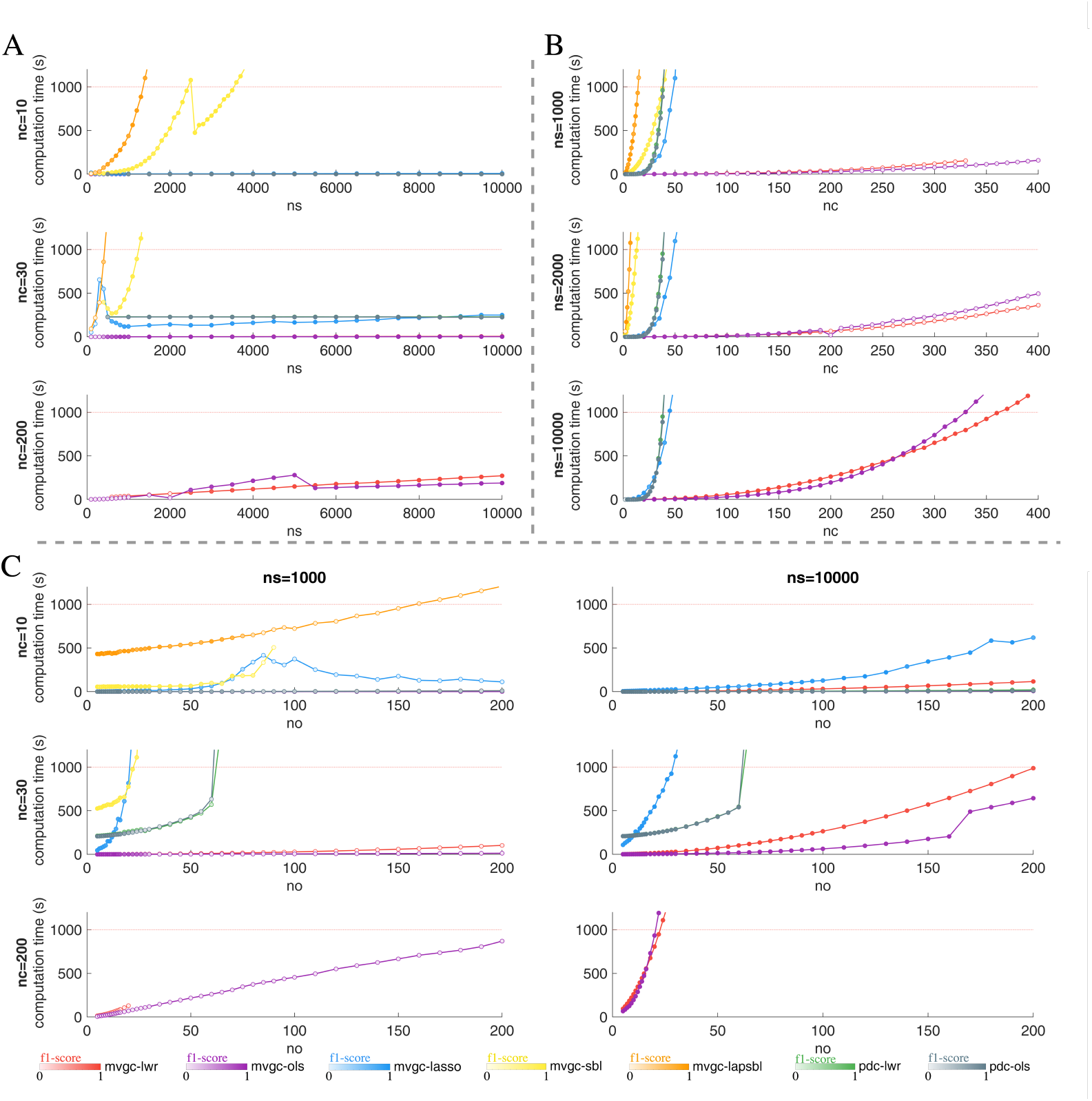
**A**: Computation time of the causality estimation with respect to the number of samples *ns*. Model order *no* is fixed to *no* = 10. From top to bottom, the number of signals is fixed to *nc* = 10, 30, 200. **B**: Computation time of the causality estimation with respect to the number of signals *nc*. Model order *no* is fixed to *no* = 10. From top to bottom, the number of samples is fixed to *ns* = 1000, 2000, 10000. **C**: Computation time of the causality estimation with respect to the MVAR model order *no*. The different frames each have a fixed combination of *ns* and *nc*. From top to bottom, the number of signals in each frame is fixed to *nc* = 10, 30, 200. From left to right, the number of samples in each frame is fixed to *ns* = 1000, 10000. **A-C**: *f* 1 *− scores* are depicted by the shade (light-low to dark-high) of the dots in each frame.

Extending upon the individual effects of each parameter on computation time, we investigate how *ns, nc, no* interact. For this purpose, all combinations of parameters are relevant in the range of interest, i.e., no parameter can be fixed. As a result, the range of *ns, nc, no* must be more targeted: *nc ∈* [5, 200], *ns ∈* [100, 5000], *no ∈* [5, 50] in steps of 5, 100, and 5 respectively. This range is sufficient to capture the effects of each parameter on computation time. For each method, we modeled the computation time (*t*(*c, s, o*)) as a function of the parameters *c* (*nc*), *s* (*ns*), *o* (*no*). As absolute computation times are dependent on implementation and hardware, we investigate their orders of magnitude. The resulting models (*m*_1_ to *m*_4_) are designed with two purposes in mind. First, identifying the nature of the interaction between the parameters (figure 5A-B). Second, to estimate the order of magnitude with which each affects computation time (figure 5C-E).

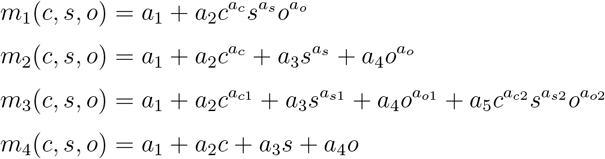

**Figure 5:**
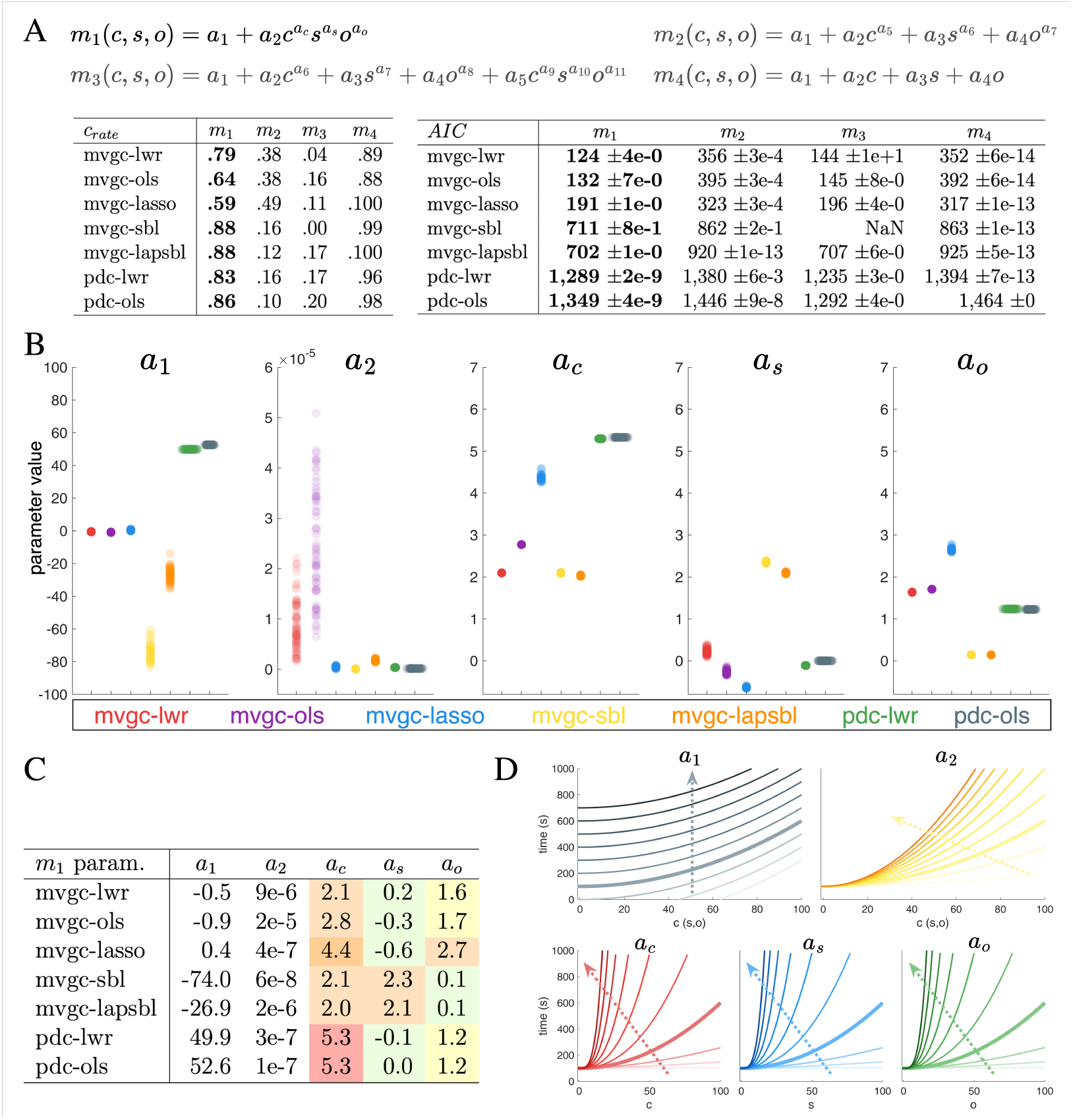
**A**: Different models to fit how computation time is affected by the number of signals (*nc*, in short *c*), the number of samples (*ns* in short *s*) and the causality order (*no*, in short *o*) in the data. Out of 100 fits with randomly drawn initial parameter values, *c*_*rate*_ indicates how many converged for each estimation method and each model. AIC indicates the average and standard deviation of the Akaike information criterion obtained by the converging fits. **B**: Distributions of the parameters of *m*_1_ from the converged fits of each method. **C**: Mean parameters of *m*_1_ from the converged fits of each method. **D**: Two-dimensional visual representation of the effect of each parameter *a*_*i*_ of *m*_1_ on computation time.

Model *m*_1_ assumes the parameter’s interaction is solely multiplicative. Model *m*_2_ assumes it is solely cumulative. Model *m*_3_ assumes both multiplicative and cumulative contributions. Model *m*_4_ serves as a sanity check that parameters affect computation time with varying powers rather than linearly. The four non-linear models are fit via Matlab’s function nlinfit, which implements the *Levenberg-Marquardt nonlinear least squares algorithm*[31] to iteratively estimate parameters *a*_*i*_ from initial values *U ∈* [0, 1]. Each model is fitted 100 times. Outliers are removed, i.e., fits for which any of its predicted parameters (*a*_*i*_) were more than three scaled median absolute deviations. To select the best-fitting model, we consider the number of fits left after the outlier removal, indicating model fit stability. We also consider the Akaike information criterion (AIC), which indicates goodness of fit while accounting for the varying numbers of parameters among the models.

### 2.4 Sparse Multivariate Granger Causality

The computational complexity of conditional Granger causality estimation is fundamentally constrained by the number of pairwise signal interactions. For *nc* signals, this results in a search space of order *O*(*nc*^2^) where *O*(*·*) denotes asymptotic upper-bound complexity. This quadratic scaling imposes a fundamental computational barrier, such that any algorithm that exhaustively explores the search space becomes intractable for large numbers of signals. While the standard approach to managing computational complexity is to restrict the number of signals considered, this strategy is ill-suited to applications involving many relevant signals. Rather than limiting signal count, we propose sparse-MVGC (sMVGC), an extension of MVGC that exploits the sparsity of pairwise signal connections by constraining the number of potential interactions per signal, rather than excluding signals entirely. This reduces the search space to *O*(*nc × d*) where *d* denotes the number of potential connections from each of the *nc* signals to the others, enabling conditional Granger causality estimation across larger sets of signals simultaneously. In practice, sMVGC requires prior knowledge of the relative likelihood of connections within the search space, which is inherently application-dependent. This prior knowledge is formalized as a binary *nc × nc* adjacency matrix, in which the *nc × d* true-valued entries designate the connections retained for estimation. The sparsity of this matrix drives the scalability of the algorithm: sparser matrices yield lower values of *d*, reducing computational cost.

Algorithm 1 is a formalization of the original MVGC, while algorithm 2 describes its adaptation to sMVGC leveraging the added prior knowledge. The function labeled `estimate_var_error` refers to the various methods described in section 2.2 to solve the optimization problem described by equation 9. Figure 6A provides a visualization of both algorithms, their inputs, and the search space they each cover. Algorithmically, MVGC consists of two main components: a full regression over all signals, and a per-signal reduced regression over all signals but the one under consideration. In sMVGC, each reduced regression is instead computed over a signal-specific subset, defined by the potential connections to that signal as specified in the prior. This entails a dedicated full regression for each subset, rather than a single full regression shared across all reduced regressions as in MVGC. Consequently, a fully connected prior would render sMVGC strictly less efficient than MVGC, as it would require *nc* full regressions of equivalent size instead of one.

**Figure 6:**
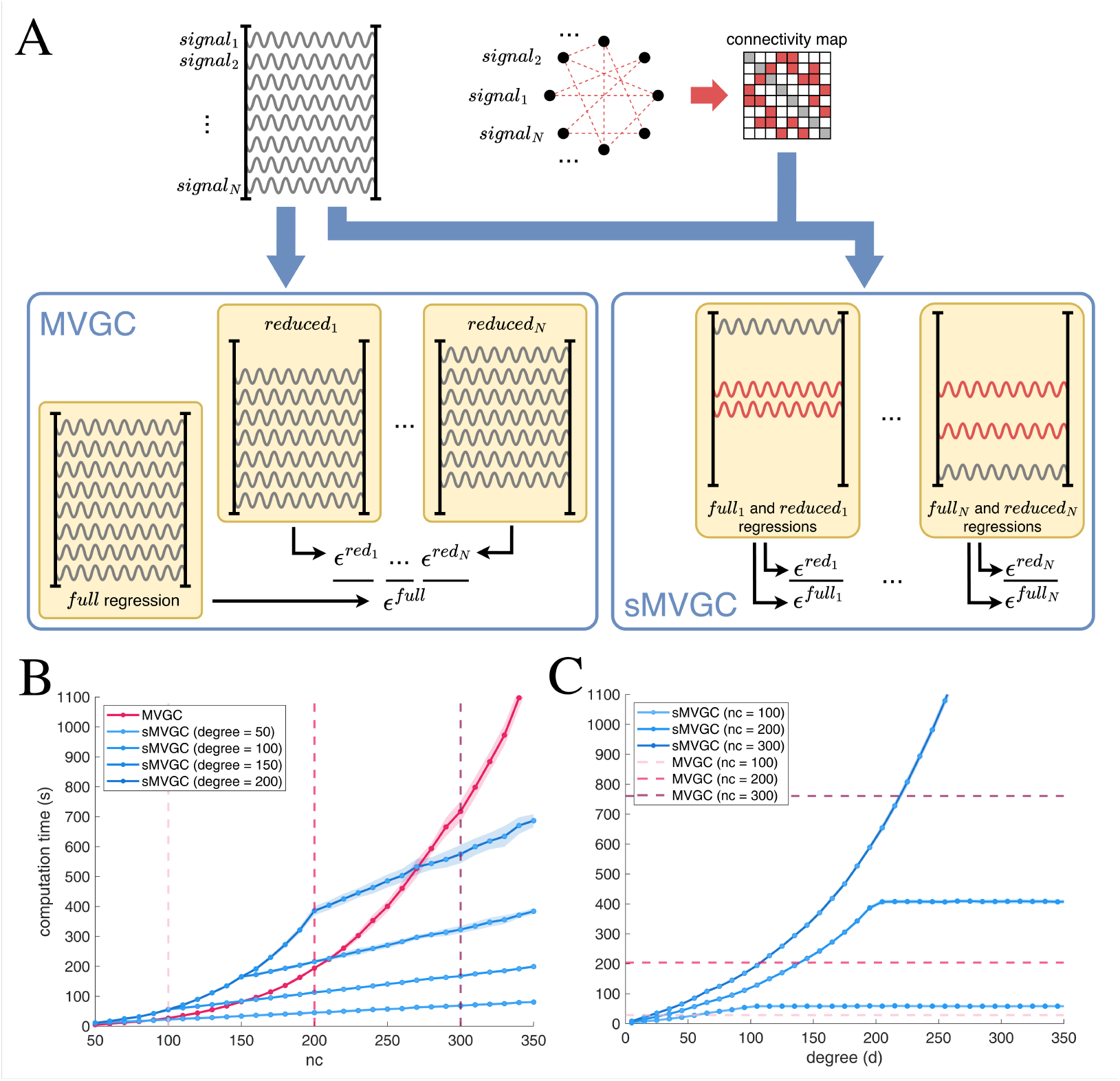
**A**: Comparative visualization of multivariate Granger causality (MVGC) with its sparse alternative (sMVGC) in terms of their inputs and inner workings. **B**: Computation time of MVGC and sMVGC with respect to the number of signals *nc*. For each *nc*, priors with varying sparsity levels (*d* = 50, 100, 150, 200) are provided to sMVGC. **C**: Computation time of sMVGC with respect to the sparsity level *d* of its prior for sets of varying numbers of signals (*nc* = 100, 200, 300).

#### Algorithm 1

MVGC (*data, order*)

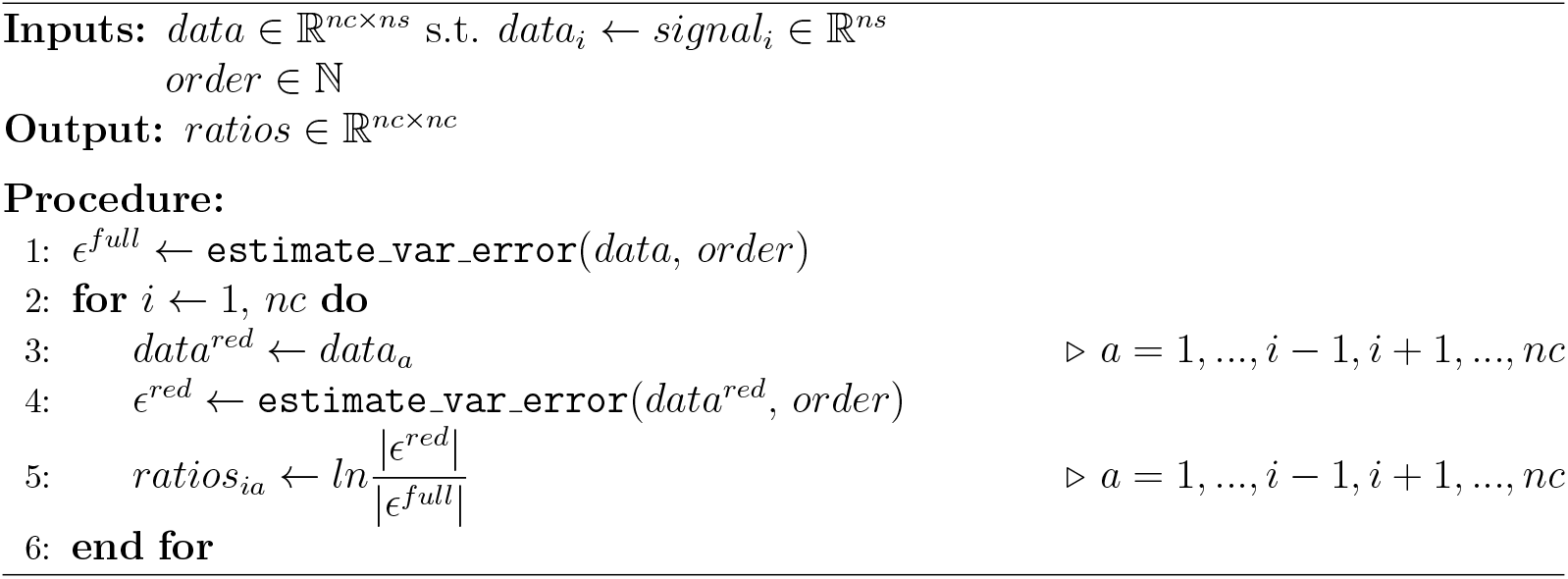

To assess the performance of sMVGC, we simulate sets of increasing numbers of signals and priors across varying sparsity levels. Granger causality is estimated using both MVGC and sMVGC; computation time is then systematically recorded for each combination of *nc* and *d*. In figure 6B, the computation times of both methods are compared with respect to the number of signals (*nc ∈* [50, 350] in steps of 10) considered simultaneously. Different levels of sparsity are considered for the prior used by sMVGC (degrees *d* = {50, 100, 150, 200}). In figure 6C, the computation times of sMVGC are displayed with respect to the sparsity of its prior (*d ∈* [5, 350] in steps of 10), and for varying numbers of signals (*nc* = {200, 300, 400}).

#### Algorithm 2

sMVGC (*data, order, prior*)

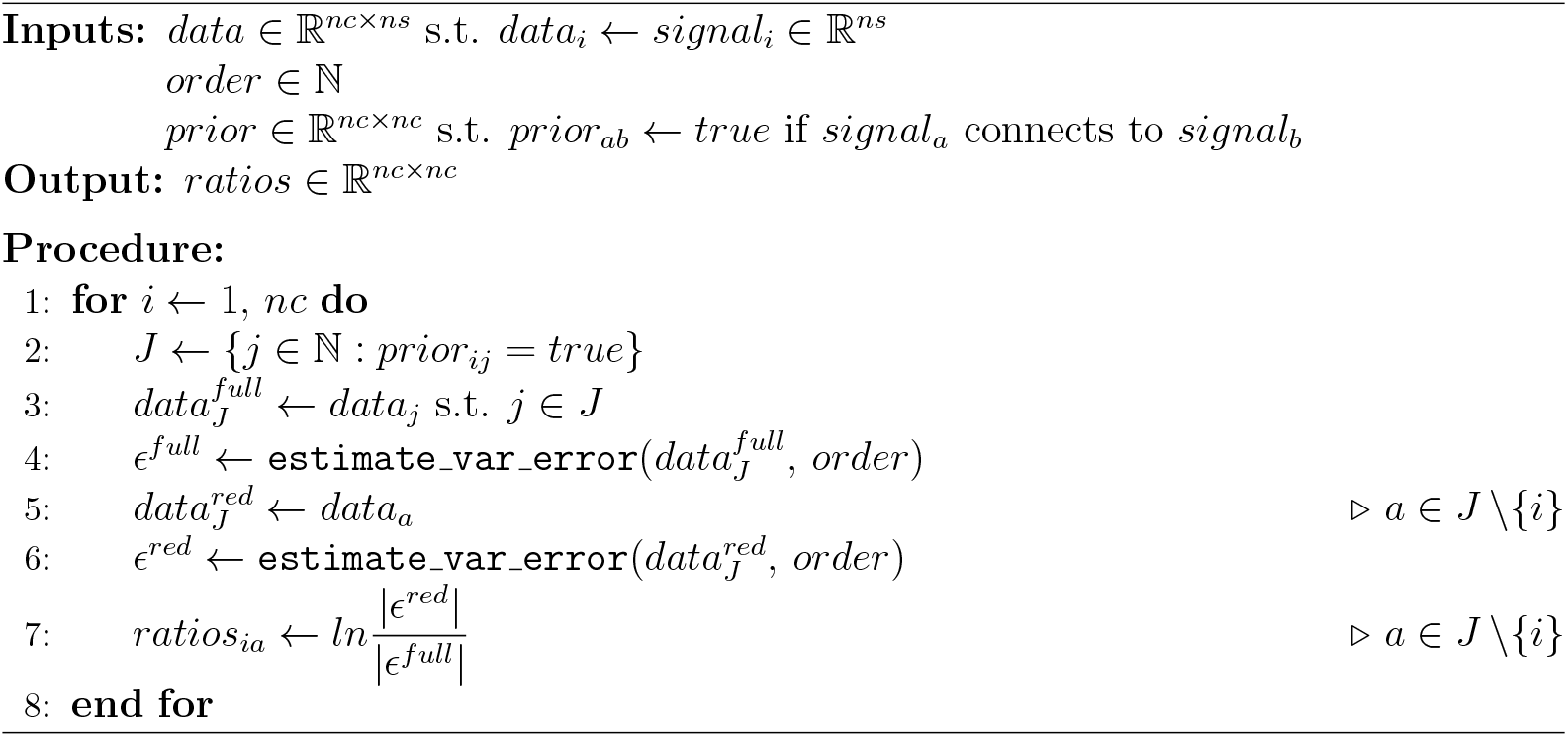

## 3 Results

### 3.1 Effect of number of signals and samples on prediction performance

Figure 2 shows that, for each method and each value of *nc*, the performance starts low, increases with *ns*, and stabilizes to high performance with high enough *ns*. This confirms that correct causal estimation depends on the length of the data (*ns*). Specifically, there seems to exist an optimal *ns*\* which is the minimum *ns* above which the prediction performance does not increase significantly. In figure 2, this optimal *ns*\* proves to depend on *nc*, i.e., predicting causality accurately between more signals requires longer recordings. This result is consistent across methods. Figure 3A allows a more precise look into this relationship between *ns* and *nc*. It depicts a *feasibility area* in which each algorithm can be used to estimate the causality with corresponding *ns* and *nc*. It is important to note that the border between the feasibility area and the black “overtime” area is driven by the arbitrary 1000 second computation time limit fixed for these experiments. Along the feasibility area, the border between light and dark green is of particular interest. It corresponds to the relation between optimal *ns* and *nc*, i.e., the function *ns*\**nc*, which is linear. Regressing the points at this border for each method yields figure 3B. Specifically, we define this border as the *ns*\* below which *f* 1*s <* 0.5 and above which *f* 1*s >* 0.5. PDC yields predictions of *f* 1*s >* 0 with much lower *ns* than the other methods, but requires a higher *ns*\* for *f* 1*s >* 0.5. The other methods have similar *ns*\* slopes around *a* = 10, indicating that 10 times more samples than signals are necessary to infer causality accurately between them.

### 3.2 Effect of number of signals, samples, and model order on computation time

In figure 4A *ns* (number of samples) affects the computation time of SBL-based methods at least quadratically. For all other methods, it does so marginally, i.e. at worst linearly, even for high *nc* (number of channels, i.e. signals). In figure 4B, *nc* affects computation time at least quadratically for all methods. Only MVGC-LWR and MVGC-OLS estimate causality correctly among hundreds of signals. Doing so accurately (dark-colored dots) requires high-enough *ns*, i.e., between 2000 and 10, 000. In both panels A and B of figure 4, the depiction of f1-scores is in line with the results in figure 2, i.e., increasing *ns* yields higher *f* 1*s*, and higher *nc* requires more *ns* to get similar *f* 1*s*. In figure 4C the effect of *no* (model order) on computation time is mostly moderate, but highly depends on the other two parameters. Regarding the *f* 1*s* with respect to *no*, it seems to be playing the same role as *nc*. With fix *nc* and *ns*, increasing *no* reduces *f* 1*s*. Increasing *ns* allows accurate estimation with higher *no*. It suggests that increasing *no* requires higher *ns* to estimate with similar accuracy. Overall, figure 4 indicates that *nc* has the most drastic effect on computation time. Additionally, all three parameters seem to interact in their effect.

### 3.3 Interaction of number of signals, samples, and model order

In figure 5B, the right table reveals that overall, model *m*_1_ has the best Akaike information criterion (AIC) values regardless of the method. Another model, *m*_3_, has similar AICs across methods but has a very low converging rate (*c*_*rate*_). The fact that, for each causality estimation method, model *m*_3_ converged in less than 20% of the fits indicates that it is less robust. As the sole model with both small *AIC* and high *c*_*rate*_, *m*_1_ is considered the best model to fit the data for each method. That *m*_1_ fits the data better than *m*_4_ confirms that *nc, ns*, and *no* affect computation time with different power scales (*a*_*c*_,*a*_*s*_,*a*_*o*_), rather than linearly. That *m*_1_ fits the data better than *m*_2_ and *m*_3_ indicates that their interaction is mostly multiplicative.

As *m*_1_ best fits the data of all methods, its learned parameters can provide insights into the effects of *nc, ns*, and *no* on causal estimation computation time. In figure 5C, the parameters of importance (*a*_*c*_, *a*_*s*_, *a*_*o*_) are less variable than they differ from one another, allowing their comparison. Figure 5D summarizes the parameter values as the mean from panel C. For most methods, *a*_*s*_ ~ 0 which indicate that *ns* hardly affects the computation time. An exception are the SBL-based methods, which scale quadratically with the number of samples *ns*. In practice, *ns* typically ranges in the thousands while *nc* and *no* range in the dozens, making *a*_*s*_ critical for the overall scalability of a method. That the *a*_*s*_ of SBL-based methods is higher than the others limits its use. This explains why the feasibility area in figure 3A is so limited for those methods. The power term of *no* is moderate, i.e. *∈*]1, 2[for most methods. It is almost zero for SBL-based methods, indicating an apparent invariability of computation time with respect to *no*. This is most likely due to the very limited feasibility area of SBL-methods, driven by its low scalability with respect to *ns*. It makes it difficult to test enough challenging values of *no* to fit its effect. Finally, *nc* has the largest effect on computation time. It is at least quadratic for all methods, cubic for MVGC-LASSO and MVGC-OLS, and even quintic for PDC.

### 3.4 Improving scalability with sparse Multivariate Granger causality

Figure 6B illustrates the computational complexity of both MVGC and sMVGC as a function of *nc*. MVGC exhibits quadratic growth, while sMVGC scales linearly with *nc* for *d < nc*. When *d ≥ nc*, the prior used by sMVGC becomes fully connected and complexity reverts to quadratic growth. Notably, under low-sparsity prior conditions, sMVGC requires longer computation time than MVGC. For instance, sMVGC with *d* = 200 takes longer to compute than MVGC for *nc <* 270, and sMVGC with *d* = 150 does so for *nc <* 210. The red dotted lines mark the computation times of MVGC at *nc* = 100, *nc* = 200 and *nc* = 300, which are carried over to figure 6C to facilitate a direct comparison of scalability across equivalent signal set sizes (*nc* = 100, 200, 300).

Figure 6C reveals that the computation time of sMVGC grows quadratically with the prior degree *d*, reaching a plateau at *d ≥ nc*. For *d < nc*, the rate of growth with respect to *d* increases with *nc*, indicating that larger signal sets require higher prior sparsity to stay tractable. The intersections between MVGC and sMVGC curves are particularly informative: MVGC (*nc* = 100) intersects sMVGC (*nc* = 100) near *d* = 60, MVGC (*nc* = 200) intersects sMVGC (*nc* = 200) near *d* = 140, while the corresponding intersection for *nc* = 300 occurs near *d* = 220. These crossover points define the sparsity threshold below which sMVGC offers a computational advantage over MVGC for a given *nc*.

## 4 Discussion

In the present study, we investigated how MVAR-based Granger causality estimation is limited by the scale of key parameters. We found that increasing the number of samples is necessary when estimating the causality between a large number of signals. We further revealed that increasing the number of considered signals increases the computation time of all investigated methods at least quadratically. Furthermore, the number of signals, the number of samples, and the order of the MVAR model affected the computation time of all methods multiplicatively, highlighting the need to consider them simultaneously when designing an experiment. Finally, we propose a variation of the most scalable method, exploiting the sparsity in large-scale signal causalities, to further improve its scalability. Through our results, we provide guidelines to dimension data for future MVAR-based causality analyses.

### Increasing the number of signals requires increasing the number of samples

The requirement for sufficiently long recordings to estimate MVAR parameters and infer causality is well established. Prior work has progressively refined this intuition into concrete guidelines, converging on a minimum of 10 *× nc × no* samples for systems of up to 30 signals [10, 11, 24], and up to 100 *× nc × no* for larger ones [12]. Our analyses complement these guidelines by incorporating practical computation times across a range of estimation methods (figures 2, 3), revealing an optimal recording length that balances estimation performance and signal duration. This optimal length increases with the number of signals considered, consistent with prior findings [10–12, 24].

### Computation time grows at least quadratically with the number of signals

While it is widely accepted that Granger causality is limited in the number of signals between which it can be inferred, an explicit limit is lacking in the literature. In fact, in previous work, inferring causality is often done between fewer than 30 signals [17, 24, 26, 32, 33]. Particularly in neuroscience, it is common to reduce the number of signals sampled to a limited number of regions of interest [12, 32–34]. Even work promoting “large scale” causal inference keeps the number of signals limited to below 50 [12, 35, 36]. More recent work has examined causality between around 100 sources [37, 38], but the practical limit remains blurry. Our results indicate that the computation time of all methods grows at least quadratically with the number of signals (figure 5), which explains that all previous work was limited in that regard. Going further, our results reveal that, while most methods would limit causal inference to fewer than 100 signals, MVGC-OLS and MVGC-LWR allow accurate causal prediction between more than 500 signals within an hour of computation time (figures 4, 5).

### Our conclusions generalize across various modeling choices

The interpretability of our conclusions is limited by the modeling choices made when designing the simulations. Notably, we simulate external noise with Gaussian white noise. External noise in electrophysiological signals is often comprised of background brain activity, which could be more realistically simulated by pink noise. Similarly, we simulated causal networks with a fixed node degree of 2. Increasing the number of signals (*nc*) in the simulated networks increases sparsity. Realistic node degrees in natural causal networks are likely to be denser, depending on the signal’s nature and sampling. Our causality network simulations exclude feedback loops by design to avoid the data saturation that can arise from such non-linearities. However, in natural systems, feedback is not uncommon. In our analyses, the order driving simulated causalities perfectly matches the order assumed by the causality estimation methods. In practical applications, this information is rarely known. While it can be estimated with different criteria [39], such an estimation requires a subtle balance between under- and overfitting of the finite signals [1]. Supplementary experiments investigating the effect of SNR on causality estimation reveal that MVAR parameters affect the detectability of causalities independently from the method employed (figure **??**). Further supplementary experiments confirm that our conclusions on scalability generalize to pink external noise, denser causal networks, including feedback causalities, and in the presence of underestimated model order (figures **??, ??**).

### Scalability is not affected by signal nonlinearities

Both our causal simulations and the methods of interest make the assumption that the causal relationships between signals are linear. In fact, standard MVAR models are linear by design. However, in natural contexts, nonlinearities can often arise [40, 41]. While this drives the development of causal estimation methods targeted at specific nonlinearities [36, 42, 43], not much is known about the exact properties of such measures. Moreover, inherently linear methods are not powerless to model and detect nonlinearities. For instance, previous work has shown that PDC can predict some causalities in nonlinear models [44, 45]. Our own supplementary results (figures **??, ??**) indicate that both PDC and MVGC can detect nonlinearities in the form of feedback loops, cosine, and quadratic causalities to some extent. More importantly, they suggest that nonlinearities do not affect the scalability or computation time of either method.

### Scalability is constrained by computational and stationarity considerations

While our experiments define the computation time limit to 1000 seconds for tractability purposes, it comes naturally that such bounds are arbitrary. Given more computation time and/or more computation power, causal analyses between even more signals become realistic. On the other hand, in natural contexts, longer signals are more likely to violate the stationarity assumption required by Granger causality [8]. Since causal estimation between more signals requires longer segments (figure 3), in turn, it risks violating the assumption. In applications with fast, transient causal dynamics, longer data segments become impractical, and a large number of signals cannot be considered simultaneously. This further emphasizes the need for clear guidelines on causal analyses and their parameter interactions.

### Causal inference methods trade-off scalability for robustness

Most recent causal inference methods were developed to infer causality more accurately, from as short segments as possible, and despite different noise conditions [14, 17, 26]. Our results indicate that their computation time scales worse with increasing MVAR parameter dimensionality than other methods. It suggests that recent methods have traded off computation complexity for robustness against low-sample data. While this is crucial for some applications, e.g., trial-based neurophysiological experiments, it greatly limits the methods’ use to large-scale data. Our experiments highlight the importance of considering dimension scalability in future causal inference methods.

### sMVGC leverages sparse connectivity to enable large-scale causality analyses

Sparse-MVGC (sMVGC) addresses the problem of scaling with the number of signals, *nc*. By assuming that not all signals interact, it reduces the number of potential pairwise connections, thereby enabling causality estimation among a larger number of signals. Specifically, given a fixed connectivity degree, sMVGC scales linearly with *nc*, though this becomes quadratic as the degree increases. This method thus constitutes an alternative to signal reduction in settings where a large number of signals are relevant but their pairwise connections are expected to be sparse.

Such settings are not uncommon. In electrophysiology, for instance, source reconstruction yields thousands of spatially distinct signals across the brain. The prevailing approach to managing this dimensionality relies on parcellating sources into regions of interest defined by an anatomical atlas. This procedure can introduce biases tied to the arbitrary choice of atlas. While the number of relevant sources is large, their anatomical interconnections are inherently sparse, being constrained by a limited number of white matter fiber tracts. Leveraging structural connectivity maps as priors for sMVGC could therefore enable causality analyses among thousands of sources, substantially extending what is currently feasible in whole-brain conditional causality analysis.

### Guidelines

We can summarize our results into the following practical guidelines for scalable multivariate causality analysis:

- Accurate causal inference among *C* signals requires at least 10*C* samples for MVGC, and 20*C* samples for PDC.
- Causal inference among hundreds of signals is possible with MVGC-OLS and MVGC-LWR methods. Requiring longer segments, they are best suited to detect long stationary causalities (stationarity assumption).
- MVGC-OLS should be favored to detect dense underlying causal networks (average node degree *>* 50) from hundreds of signals.
- In situations where thousands of signals (*nc*) are relevant, but their interconnections are expected to be sparse, sMVGC is the only reasonable option.
- More data-efficient methods, such as LASSO, SBL, and lap-SBL, can infer causality among dozens of signals at most, within a reasonable time. Requiring shorter segments, they are best suited to detect faster transient causalities (stationarity assumption).
- Overestimating data order provides better causal detection than underestimating it, at the cost of increased computation times.

In conclusion, our findings provide a clear practical range for the number of signals, samples, and model order to infer multivariate Granger causality with current methods. We provide guidelines to dimension future causality analyses. We propose sMVGC, an adaptation of multivariate Granger causality that exploits sparsity in pairwise signal connectivity to enable analyses at a scale beyond that of existing methods. Our results can help select an adequate MVAR-based causal estimation method for the size of the data to analyze. More fundamentally, this study highlights the importance of considering dimensional scalability in Granger causality analyses, as well as future method developments.

## Data availability

All resources necessary for the replication of this study are made available online. The source code can be found in a public GitHub repository^1^. The source code allows generating all the data relevant to this research. Generating it all can require weeks of computation time. Alternatively, the resulting data can be found on the Open Science Framework^2^.

### Glossary

AIC: Akaike information criterion.
EM: expectation-maximization algorithm.
FN: false negatives.
FP: false positives.
gPDC: generalized partial directed coherence.
lapSBL: Laplace sparse bayesian learning.
LASSO: least absolute shrinkage and selection operator.
LWR: Levinson-Wiggins-Robinson regression.
MVAR: multivariate autoregressive models.
MVGC: multivariate Granger causality.
sMVGC: sparse multivariate Granger causality.
nc: number of signals (channels).
no: MVAR model order.
ns: number of samples.
OLS: ordinary least-squares regression.
PDC: partial directed coherence.
SBL: sparse bayesian learning.
SNR: signal-to-noise ratio.
TN: true negatives.
TP: true positives.

## Author contributions

**Thomas Pirenne**: Conceptualization, Formal Analysis, Investigation, Methodology, Software, Writing – Original Draft Preparation. **Esther Florin**: Conceptualization, Funding Acquisition, Project Administration, Supervision, Writing – Review & Editing.

## Acknowledgments

We would like to thank the members of the Institute of Clinical Neuroscience and Medical Psychology of Heinrich Heine University and in particular Fabrice Hubschmid and Oliver Kohl for their helpful feedback.

## Footnotes

1 https://github.com/TPirenne/granger-causality-scalability

2 https://osf.io/ws2ph/

