## Supplementary Materials for "Parameter scaling of multivariate Granger causality"

### Appendices

#### A Effective coupling strength

##### A.1 Experiment

The experiments listed in section 2 of the main manuscript focus on scalability. The analyses described in its section 2.3 were performed without external noise. We ran them again with an SNR=1 to verify whether external noise affects scalability. Moreover, external noise affects the effective coupling strength introduced in section 2.1 as per equation 3. It provides fine control over the effective coupling strength in our simulations, which, in turn, allows us to investigate the extent to which the effective coupling strength affects causal inference by our methods of interest. We simulate 100 trials of fix and reasonable $nc$ (15), $ns$ (500), and $no$ (10). Distributions of coupling strengths are spread uniformly between -1 and 1 ($\mathcal{U} \in [-1.0, 1.0]$). For each method and for $SNR = \{1/3, 1, 3\}$, we estimate causality from the generated times series. We then discriminate detected (True Positive) and missed (False Negative) connections and identify their corresponding coupling strengths from the simulation. We then train a simple binary classifier (binary decision tree with 2 splits - Matlab's `fitctree`) to derive a coupling strength threshold separating TP from FN. The resulting threshold is the *coupling strength detectability threshold*, above which most causal connections are detected and below which most are not. In a subsequent experiment, we estimated this detectability threshold for combinations of parameters of increasing $nc$ $(8 - 20)$, $ns$ $(300 - 5000)$, $no$ $(5 - 20)$. Observing how the detectability threshold is affected by SNR, estimation methods, and parameters $nc$, $ns$, $no$, explains how external noise affects causality prediction performance.

#### A.2 Results: detectability threshold

Figures 2-5 were derived from analyses based on data simulated without external noise. To understand the effect of noise, all analyses have also been run with SNR = 1. Interestingly, SNR hardly affects computation time and, as such, scalability. However, it does affect correct causality estimation. Although it is evident that noise would hinder the estimation, the mechanism by which it does is of interest. The proportion of noise in the data effectively reduces the coupling strength between its signals, to a point where the causalities become undetectable. The effective coupling strength (introduced in equation 3 as $S^{eff}$) below which a causality becomes undetectable is the *detectability threshold*. Our simulation environment allows controlling the coupling strength, which makes it possible to estimate this threshold.

In Figure A.1, we display estimates of the detectability threshold for each method of interest and with different SNRs. The thresholds of all methods is around $S^{eff} = 0.4$ in noiseless conditions. These thresholds are lowered to around $S^{eff} \simeq 0.3$ for $SNR = 1$. Overall, the detectability thresholds indicate that there is an effective coupling strength above which causalities can be detected by a given method. Increasing SNR reduces the effective coupling strength, pushing a higher proportion of causalities below the detectability threshold and, in turn, reducing prediction performance.

In Figure A.1, estimated detectability thresholds prove to be dependent on parameters *ns* and *no*. For fix SNR (1/3), *nc* (15) and *no* (10), increasing *ns* lowers the detectability threshold. Fixing SNR implies that the effective coupling strengths remain unchanged. In such a situation, a reduction in the detectability threshold implies greater sensitivity to causalities, as more evidence is accumulated over longer signals. The case of fix SNR (1/3), *nc* (15) and *ns* (500) with increasing *no* is the opposite situation, yielding increasing detectability thresholds. Increasing *no* makes causal estimation more complex, while fixing *ns* limits the evidence accumulation, reducing the sensitivity to causalities as a result.

### B Simulation validation

#### B.1 Simulation design does not affect computation time

Figure B.1 shows how the computation time of methods of interest to estimate causality grows with the number of channels, with respect to different simulation design choices. Neither the external Gaussian nor pink noise affects the computation time of any method. It remains unchanged whether the causalities simulated are linear or include nonlinearities such as feedback loops, cosines, or quadratic connections. Most methods have no effect on the causality network density or computation time. One notable exception is MVGC-LASSO, which requires longer computation time to detect denser causal networks.

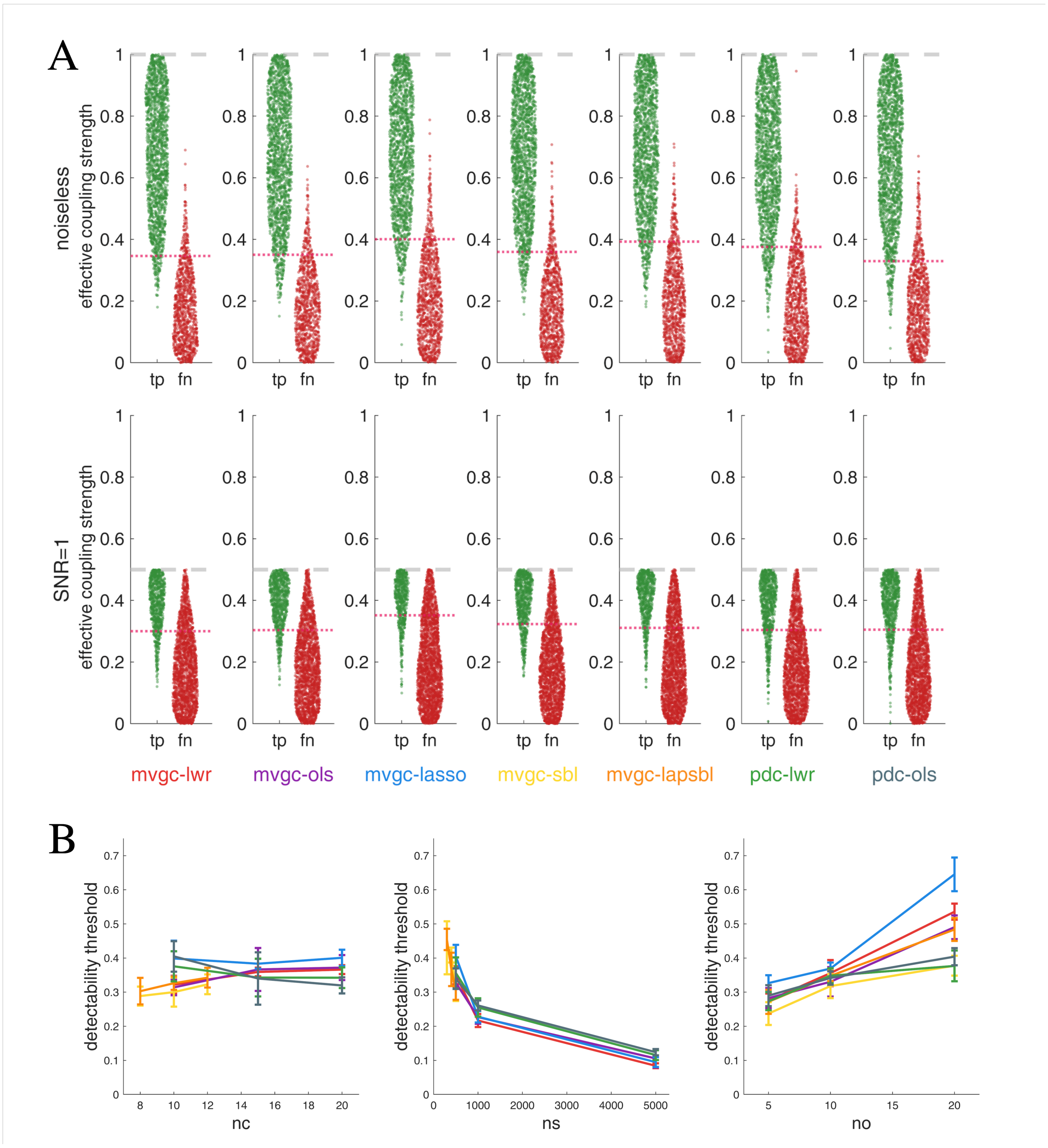


Figure A.1: **A:** True positive (tp) and false negative (fn) distributions of causal predictions with respect to the effective coupling strength of their corresponding connection. True positives are detected causalities, and false negatives are missed causalities. Red dotted lines indicate the *detectability threshold* estimated from the distributions. Grey dotted lines represent the proportion of signal with respect to noise in the simulated data. **B:** Detectability thresholds estimated from distributions of causalities predicted with varying numbers of signals, samples and order. Whiskers illustrate the standard deviation of each detectability threshold.

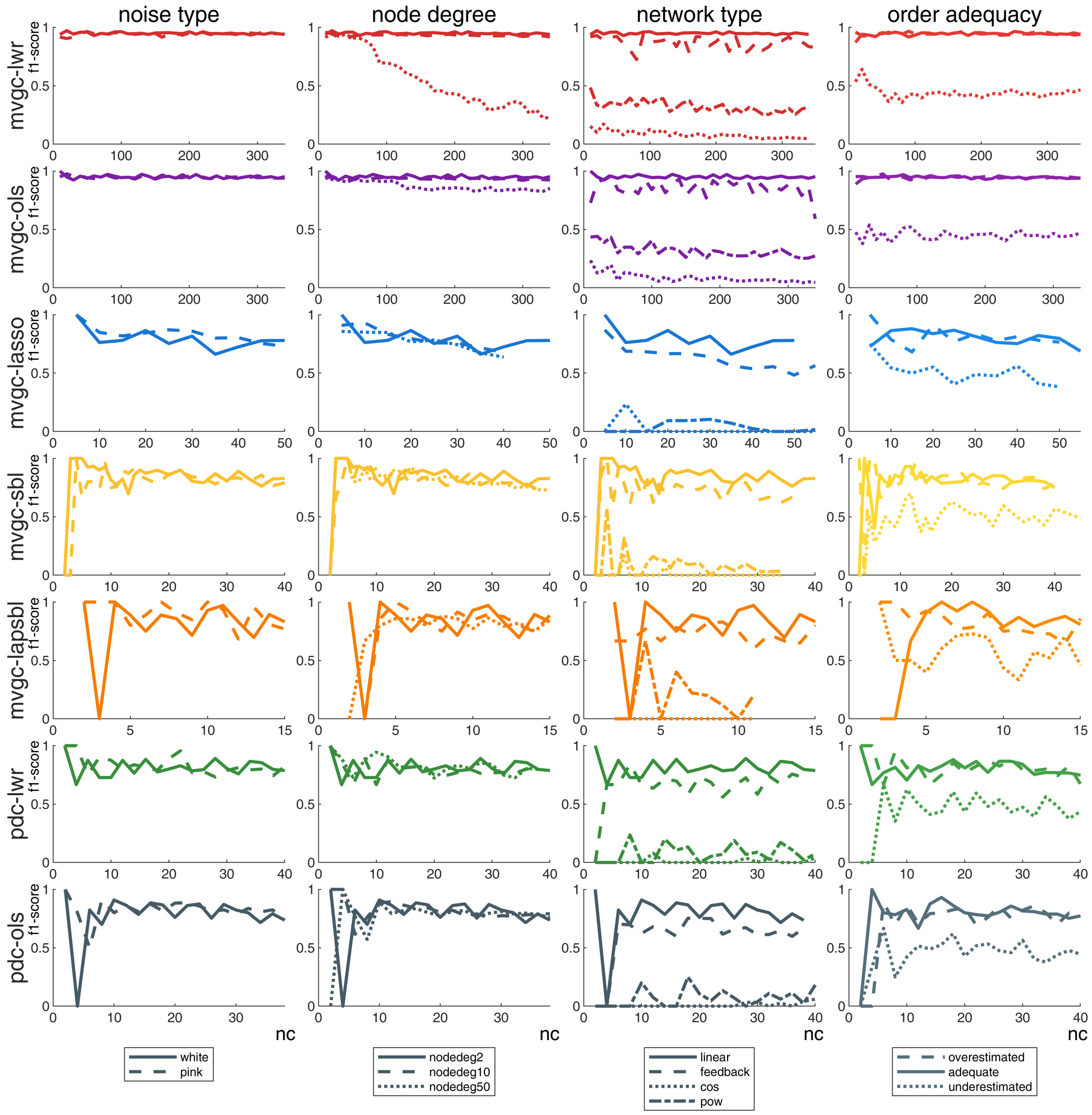


Figure B.1: Causality estimation computation time with respect to the number of signals simultaneously considered by different methods of interest (top-to-bottom). Each set of computation times is evaluated in different external noise conditions (left), with varying causal node degrees (central-left), in conditions with different types of causal relationships (central-right), and with varying data-to-estimation order mismatches (right).

#### B.2 External noise type does not affect causality estimation performance

The leftmost column of figure B.2 indicates that pink external noise scarcely affects causal estimation performance. Because pink noise is self-modulated, it is possible that such noise could lead the methods to detect spurious self-causalities. However, these insights fall out of scope, since the methods of interest in our experiments are not designed to detect self-causality.

#### B.3 Causal network density affects causality estimation performance between large numbers of signals

The central-left column of figure B.2 reveals that causal node degrees do not affect causal estimation performance between relatively low numbers of signals ($nc > 100$). However, *mvgc-lwr* and *mvgc-ols* can provide estimates of causalities among more signals in a reasonable time. Their results indicate that higher node degrees can reduce the performance of causality estimation at high $nc$. The higher $nc$, the more it seems to be reduced. A fixed node degree among increasing numbers of signals implies increasing amounts of causalities to detect, which could explain these results in $mvgc - lwr$. The method $mvgc - ols$ has this behavior to a much lower extent.

#### B.4 Causality linearity affects estimation performance but not scalability

The multivariate autoregressive model at the source of all considered methods assumes linearity of causalities. As such, fitting non-linear causalities based on feedback, cosine, and quadratic relationships is limited to linear approximations. Figure B.2 shows that feedback can be estimated with accuracy comparable to the linear case, while cosine and quadratic relationships yield much lower performance ($f1s < 0.5$). Causality prediction performance across all network types appears independent of the number of signals ($nc$) considered.

#### B.5 Model order underestimation affects causality estimation performance but not scalability

In the rightmost column of figure B.2, the overestimated and adequate model order show similar f1-scores $\sim 1$, while the underestimated models reach at most $f1score = 0.5$. It suggests that overestimating the order in causal models does not hinder prediction. Still, overestimating the model order ($no$) increases computation time, as established in subsection 3.2. More importantly, underestimating the model order affects prediction

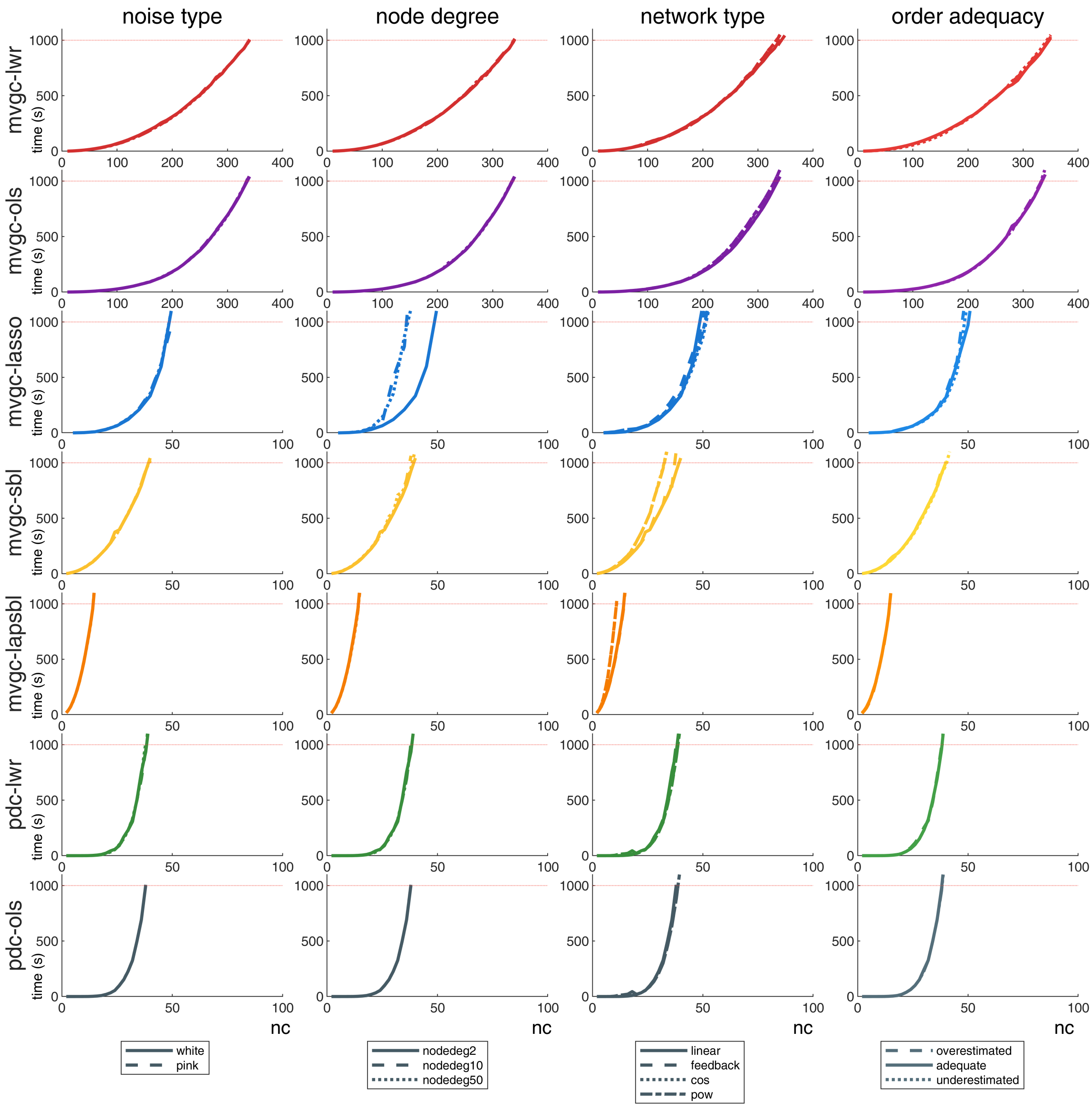


Figure B.2: Causality estimation performances with respect to the number of signals simultaneously considered by different methods of interest (top-to-bottom). Each set of estimation performances is evaluated in different external noise conditions (left), with varying causal node degrees (central-left), in conditions with different types of causal relationships (central-right), and with varying data-to-estimation order mismatches (right).

performance. Specifically, all causalities with delays greater than the model order cannot be detected. In the underestimated case in figure B.2, delays are assigned randomly following a uniform distribution between $[1, 20]$ while model order is $no = 10$. In such a condition, causalities with delays $\in [11, 20]$ cannot be detected, which represents $\sim 50\%$ of all causalities, and explains that the f1-score of such a model tops at 0.5.
